# GIS-based approach and multivariate statistical analysis for identifying sources of heavy metals in marine sediments from the coast of Hong Kong

**DOI:** 10.1101/2022.07.16.490032

**Authors:** Fengwen Huang, Chen Chen

**Affiliations:** Key Laboratory of Neuroscience, Department of Biomedical Science, City University of Hong Kong, Hong Kong; Shenzhen Key Laboratory of Marine Bioresources and Ecology, College of Life Sciences and Oceanography, Shenzhen University, Shenzhen, China

**Author notes:** The first author in this paper. Corresponding author. *E-mail addresses*.

**Keywords:** Heavy metals, Sediment, Source identification, Statistical analysis

## Abstract

Multiple methods consisting of geographic information system (GIS) technique, enrichment factor (EF), potential ecological risk index (PEI) and multivariate statistical methods was developed to identify anthropogenic heavy metal sources in marine sediments of Hong Kong. The distributions of heavy metals in sediments have been analyzed, and their pollution degrees, corresponding potential ecological risks and source identifications have been studied using geo-accumulation index, potential ecological risk index and integrated multivariate statistical methods, respectively. Three different types of anthropogenic inputs could be identified via multivariate analysis. Acoording to the findings, the first principal component might originate from the industrial discharges and shipping activities. The second principal component were identified from the natural sources. The third component mainly from the municipal discharges and industrial wastewater. These results provide baseline information for both the coastal environment management and the worldwide heavy metal distribution and assessment.

## 1 Introduction

The widespread contamination by persistent and prevalent pollutants is now a major concern for the public, because of their toxicity, persistence, and bioaccumulation properties (El-Shahawi et al., 2010; Mrema et al., 2013; Ali et al., 2019). In marine environments, persistent pollutants could be further removed rapidly from the seawater by the process of sedimentation, which may directly result in higher levels of hazardous substances in sediment than in water bodies (Rousseau et al., 2015; Badruzzaman et al., 2019). Surficial sediment contamination as a consequence of intensive anthropogenic activities has become a recent concern (Christophoridis et al., 2019). The main contamination sources of heavy metals in the marine ecosystem originate from various pathways such as industrial sewage, agricultural effluent and municipal wastewater drainage (Sultana et al., 2015; Belhaj et al., 2016). The water solubility of all heavy metals was low in the aquatic system, which led to a serious adsorbed and accumulated on bottom sediments (Namieśnik et al., 2010; Wu et al., 2020). While settled metals in sediments may be re-suspended and caused secondary contamination to the aquatic environment, due to sediments act both as a sink and a source for metals in the polluted water bodies (Hiller et al., 2010; Selbig et al., 2013). This process converts the sediments into a permanent record of anthropogenic pollutant inputs. Therefore, spatial surveys of heavy metal concentrations in the coastal sediments and then comparisons with non-polluted baselines are necessary to understand the mechanisms of accumulation and geochemical distribution of heavy metals in the aquatic systems and provide basic information for the assessment of environmental risks.

To date, various methodologies have been used to assess the ecological risks of heavy metals. However, most methods are suitable only for ecological assessment of a single pollutant (e.g., Enrichment factor analysis or Geoaccumulation index method). In actual fact, multiple heavy metals usually accumulate simultaneously and cause combined pollution in sediments. Hakanson developed the potential ecological risk index to address this issue, which introduced a toxic-response factor for a given substance (Hakanson, 1980). This approach can be used to assess the combined pollution risk of multiple types of heavy metals to the ecological system. Additionally, multivariate data statistical is a proper supplementary method to identify anthropogenic contamination profiles and possible sources in a highly polluted environment (Dong et al., 2018; Zhang et al., 2020; Passarella et al., 2022). For instance, principal components analysis (PCA) simplifies the visualization of complex data sets for exploratory analysis and is used to determine the relationship between the contaminants in the sediment and their potential sources (Du et al., 2019; Camacho et al., 2020).

In the present study, sediment quality in Hong Kong marine is assessed through a data matrix, which includes ten parameters from the 2018 monitoring program. Pollution indicators were used to find out the main persistent pollutants and pollution hot spots in Hong Kong marine. The integrated application of multivariate statistical models and GIS was introduced to distinguish anthropogenic heavy metals in marine sediment of entire Hong Kong-based on datasets. The present study aimed to: (1) provide the concentration and distribution of some heavy metals in the sediments from the coastal areas of Hong kong by employing the GIS mapping technique. (2) evaluate some heavy metals’ potential ecological risk levels by applying the potential risk index method. (3) identify the sources of the heavy metals with multivariate analyses. Significantly, this study will assist the concerning authorities related to offshore conservation and administration to carry out necessary steps for planning proper management of the marine environment.

## 2. Material and methods

### 2.1 Study setting and data collection

Hong Kong is located in southern China (22°90′-22°37′N, 113°52′-114°30′E), with a land area of 1104 km^2^ and 1651 km^2^ of marine waters (Gu et al., 2019). As one of the most densely populated areas globally, with approximately 7.3 million population predominantly living along the coastal fringe of northern Hong Kong Island and the southern Kowloon Peninsula (Wang, 2020). Owing to rapid urbanization and industrialization in the 1970s, the local coastal environment has experienced unprecedented marine ecosystem pollution (Wang et al., 2013; Gu, 2019). Although effective management policies have been issued to control marine ecosystem contamination, heavy metals in the coastal zone still pose serious potential ecological risks to aquatic organisms and human health.

Bottom sediment samples in the fine-grained fraction (<63 μm) were collected by taking the top 10-cm layer of marine sediment using a grab sampler [Hong Kong Environmental Protection Department (HKEPD)]. The marine sediment samples were analyzed by HKEPD’s laboratories and the government laboratory following standard methods (American Public Health Association 1995; American Society for Testing and Materials 2001). For this study, heavy sedimentary metal and fine-grain-size data(< 63 μm) were downloaded from the open database for Marine Water Quality Monitoring Data at the HKEPD website (https://www.epd.gov.hk/epd/epic/english/datamarine.html). The selected datasets contain information on ten heavy metals monitored twice annually at 59 stations (Fig. 1) in 2018. Table 1 listed the descriptive statistics for these ten heavy metals.

**Fig 1.**
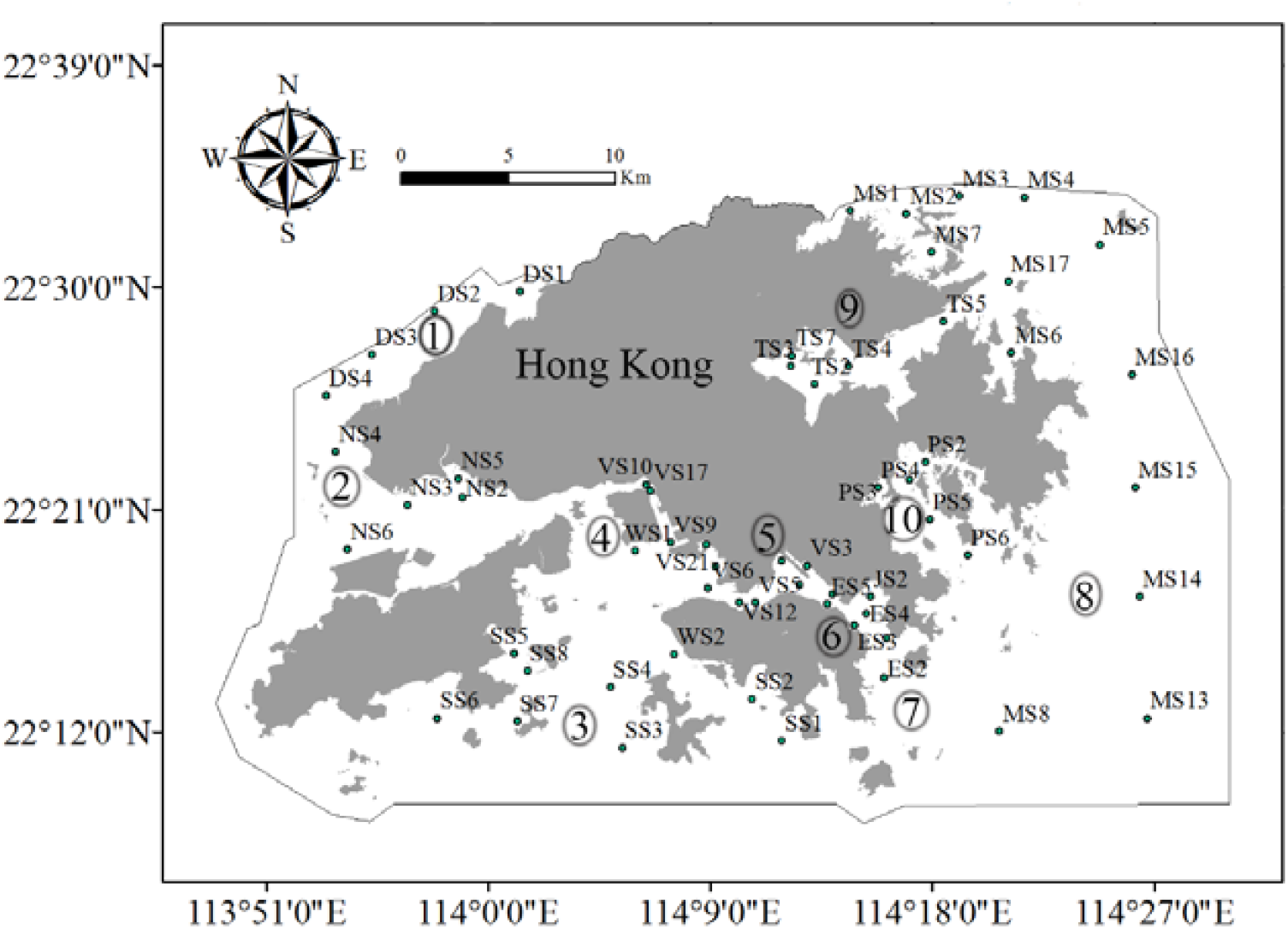
Area of study with the locations of monitoring sites (♦) and the boundary of Hongkong (—); 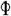: Deep Bay; 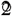: North Southern Bay; 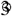: Southern Bay; 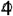: Western Buffer; 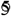: Victoria Harbour; 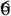: Junk Bay; 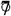: Eastern Buffer; 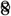: Mris Bay; 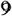: Tolo Harbour; 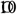 Port Shelter.

**Table 1.**
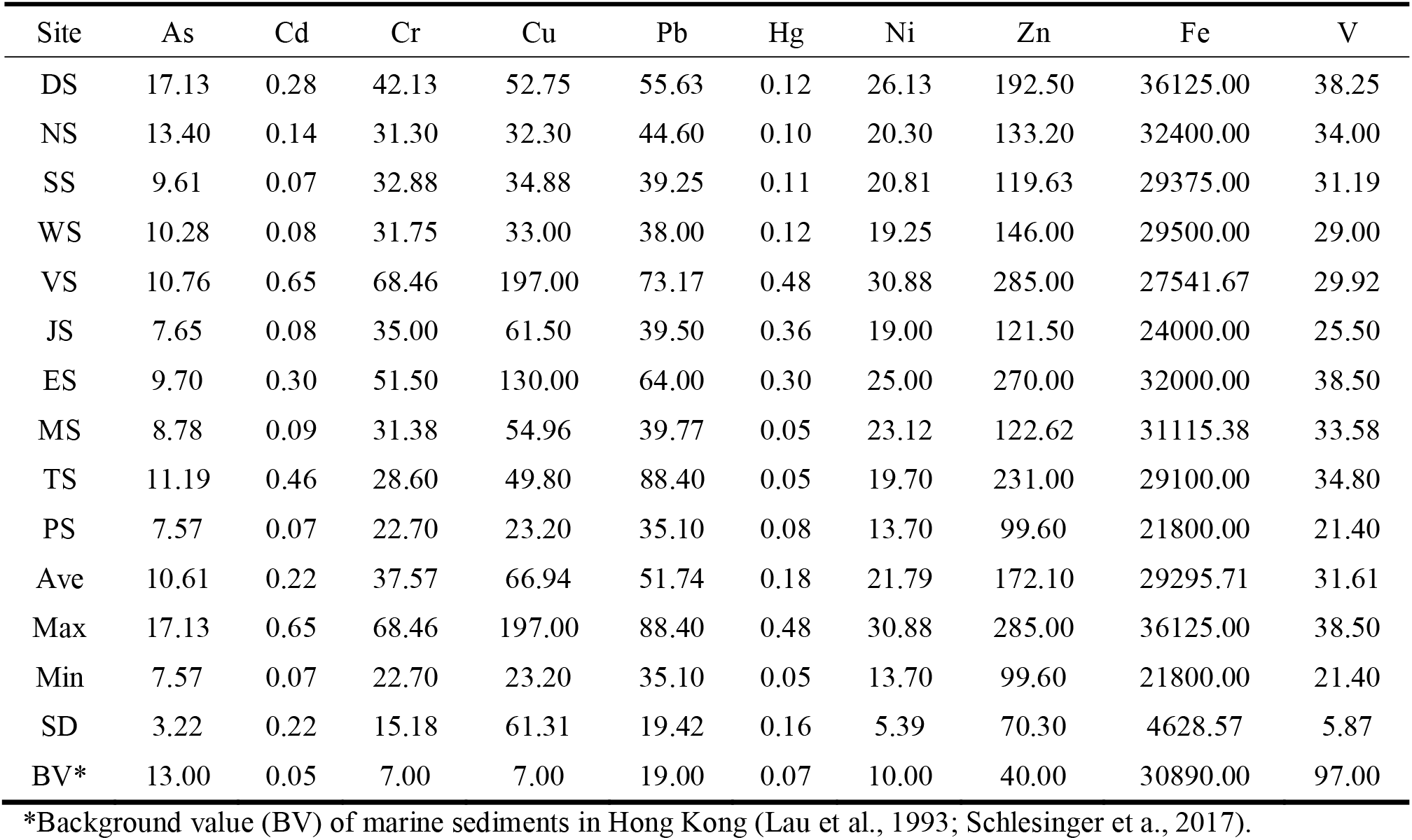
Basic statistics of heavy metal concentrations in Hong Kong’s marine sediments (mg/kg, dry weight).

### 2.2 GIS analysis

For this study, The inverse distance weighted (IDW) approach using ArcGIS 10.2 software (ESRI Inc., Redlands, CA, USA) was employed to analyze the spatial distribution characteristics of heavy metals in the sediments.

### 2.3 Enrichment factor analysis

The enrichment factor (EF) is a helpful tool for determining the degree of anthropogenic heavy metal pollution. The *EF* is calculated using the relationship below:

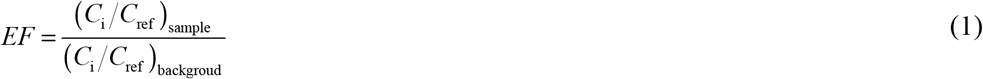

where *C*_i_ is the concentration of the element of interest and *C*_ref_ is a reference value for metals. For this study, Fe was used as the reference element for geochemical normalization for the following reasons: (1) Fe is associated with fine solid surfaces; (2) its geochemistry is similar to that of many trace metals; and (3) its natural sediment concentration tends to be uniform. Generally, *EF* values were interpreted as suggested by Sakan et al (2009), where *EF* < 2, depletion to minimum enrichment; 2 ≤ *EF* < 5, moderate enrichment; 5 ≤ *EF* < 20, significant enrichment; 20 ≤ *EF* < 40, very high enrichment; and *EF* > 40, extremely high enrichment.

### 2.4 Potential ecological risk assessment

In the present study, Hakanson’s ecological risk method is used to evaluate the potential ecological risk of metal contaminants in sediments. The ecological factor 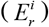 and potential ecological risk index (*RI*) of heavy metals were calculated by this method.

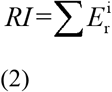

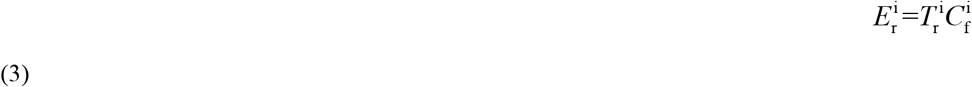

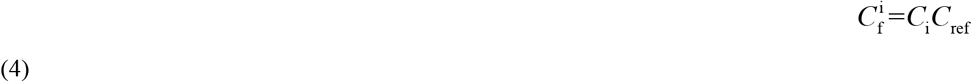

where *RI* is calculated as the sum of all risk factors for heavy metals in sediments, 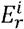 is the monomial potential ecological risk factor, 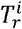 is the toxic-response factor for a given substance, which accounts for the toxic requirement and the sensitivity requirement. 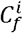 is the contamination factor, 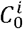 is the concentration of metals in the sediment, and 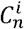 is a reference value for metals. *RI* is the sum of all risk factors for heavy metals in sediments. The relation between evaluation indices, pollution degree, and potential ecological risk are shown in Table 2.

**Table 2.**
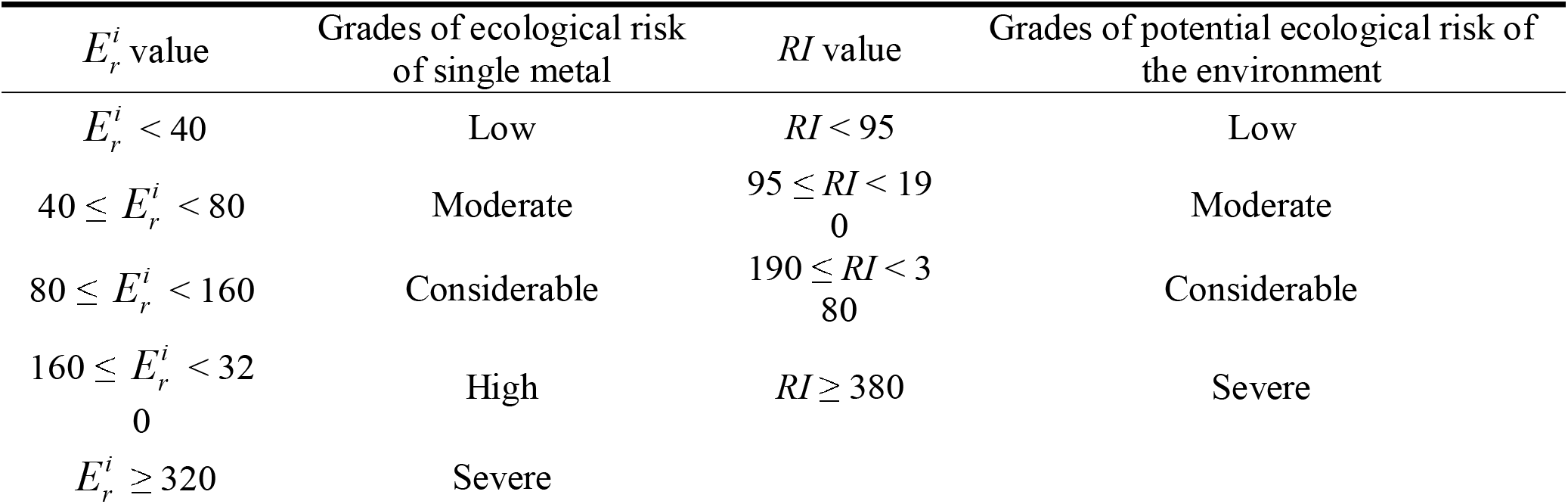
Terminology is used to describe the risk factor 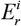 and *RI* as suggested by Hakanson (1980).

### 2.5 Statistical analysis

Multivariate statistical analysis methods such as PCA and Cluster analysis were used to determine the relationship between the contaminants in the sediment and their potential sources. The statistical software IBD-SPSS (SPSS Inc., Chicago, IL, USA) was employed in this present study. The data was graphically presented using Sigma plot 12.5 (Systat Software Inc.).

## 3. Results and Discussion

### 3.1 Distribution of heavy metals in study areas

The distributions of metal concentrations within the study area can be displayed visibly by the inverse distance weighted technique of Arc GIS. The spatial distribution maps of the ten elements are shown in Fig. 2. The element As, V and Fe displayed similar distribution patterns with relatively low concentrations in the study area. The mean concentrations of As (10.61 mg/kg) were comparable to the background value (13.00 mg/kg), while both the average concentrations of Fe (29295.71 mg/kg) and V (31.61 mg/kg) did not exceed the background value of the two elements (30890.00 mg/kg for Fe; 98.00 mg/kg for V). Indicating that the three elements in sediments in the study areas were present at unpolluted levels. Furthermore, the spatial distribution patterns of Ni and Cr were similar, and not significantly different within the study areas. The concentrations of the two elements were higher in the DS and WS sites. And relatively lower concentrations in other areas. Moreover, both the average concentrations of Ni (172.10 mg/kg) and Cr (37.57 mg/kg) were significantly above their background values (10.00 mg/kg for Ni; 7.00 mg/kg for Cr). These results suggest that the distributions of Ni and Cr in the study areas are affected by anthropogenic sources. Noticeably, the Hg showed quite different spatial distribution patterns that the higher concentrations of Hg mainly existed in the VS and ES, but lower concentrations in other areas. The average concentration of Hg (0.17 mg/kg) in all sites exhibited remarkable higher than that in the background (0.07 mg/kg). This data implies that the pollution of Hg in the study areas might be derived from the coastal point source.

**Fig 2.**
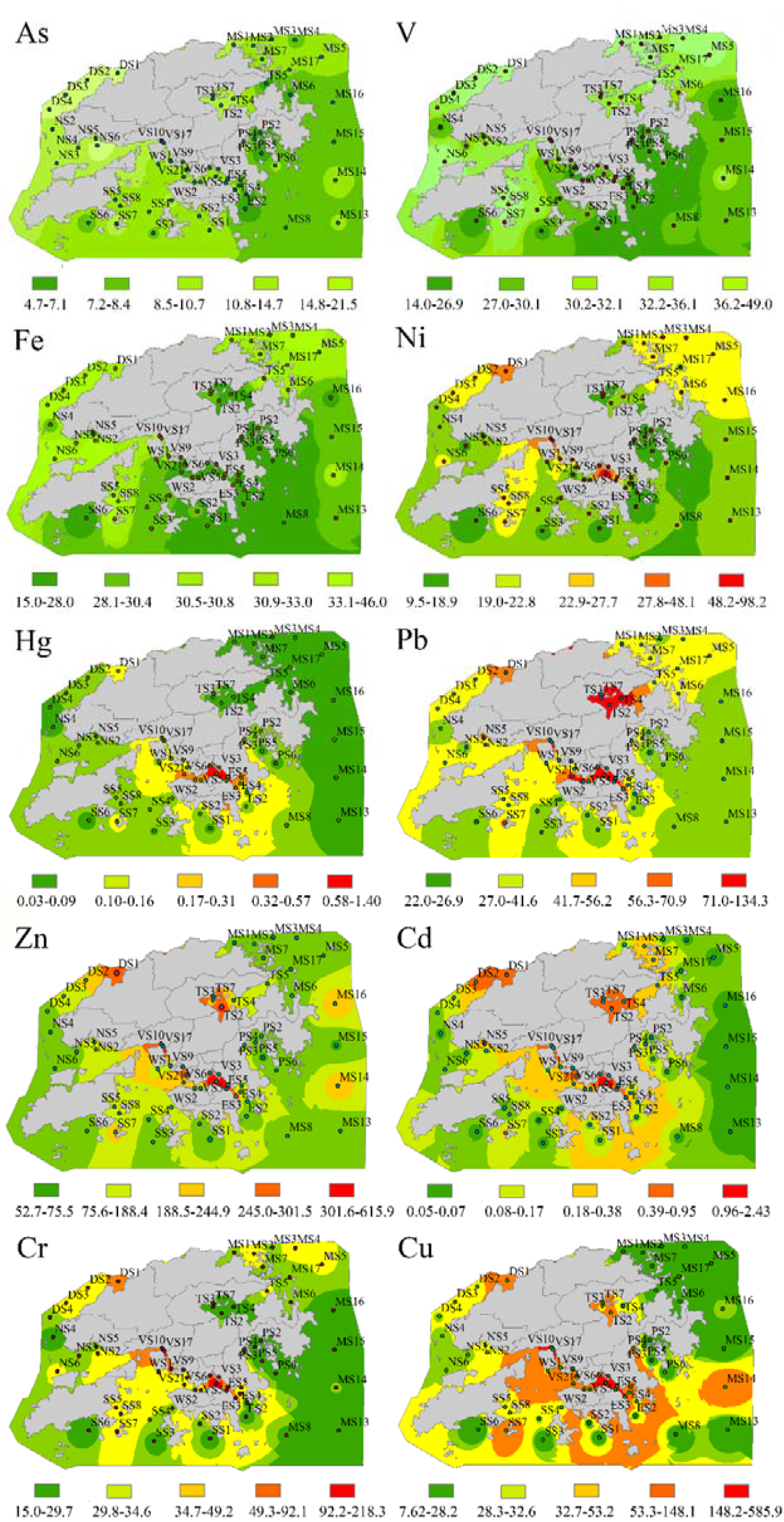
Spatial distribution of heavy metals in the coastal area in Hongkong.

In addition, we found that the Pb, Zn, Cd and Cu manifested almost identical spatial distribution patterns with higher concentrations in most of the coastal zones. The average level of four elements in the sediments (66.94 mg/kg for Cu; 0.22 mg/kg for Cd; 172.10 mg/kg for Zn; 51.74 mg/kg for Pb) were bigger than that in background (7.00 mg/kg for Cu; 0.05 mg/kg for Cd; 40.00 mg/kg for Zn; 19.00 mg/kg for Pb). Implying that the sediments in the coastal areas were seriously contaminated by these heavy metals. Overall, the spatial distribution of trace elements revealed decreasing trend from the inner estuary to the outer estuary.

### 3.2 Enrichment factor analysis

The *EF* is a convenient measure of geochemical trends and is applied for speculating on the lithogenic or anthropogenic origin of heavy metals (Wong et al., 2017). It has been postulated that a value of *EF*□≤ 2 indicates that heavy metal could result from crustal sources or natural weathering processes. The *EF* value between 2 and 5 suggests that a considerable portion of the heavy metals are delivered from non-crustal materials and the enrichment is mainly due to anthropogenic inputs. While EF value above 5 implies severe contamination by the anthropogenic sources. Therefore, the average enrichment factors of all elements were computed and presented in Fig.3. The decreasing trend of average contents’ enrichment factors was Cu > Cr > Cd > Zn > Pb > Hg > Ni > Fe >As > V. When applying the foregoing contamination categories, we found that the heavy metals of As, Fe, and V in the sample sites may be as a result of crustal materials or natural weathering processes. In contrast, the Ni, Hg, Zn, Pb and Cd having EF values between 2-5 in the sediment samples, suggesting that the contamination of these metals could be correlated to the anthropogenic sources. Nevertheless, the EF values of Cr and Cu were higher than 5 in the sediment samples, indicating that these heavy metals may be enriched due to being seriously polluted by anthropogenic inputs. Generally, it can be speculated that these metals of the studied area (Except Fe, As and V) originated from anthropogenic sources, revealing that heavy metals contaminated by human activities might have occurred and deteriorated.

**Fig. 3.**
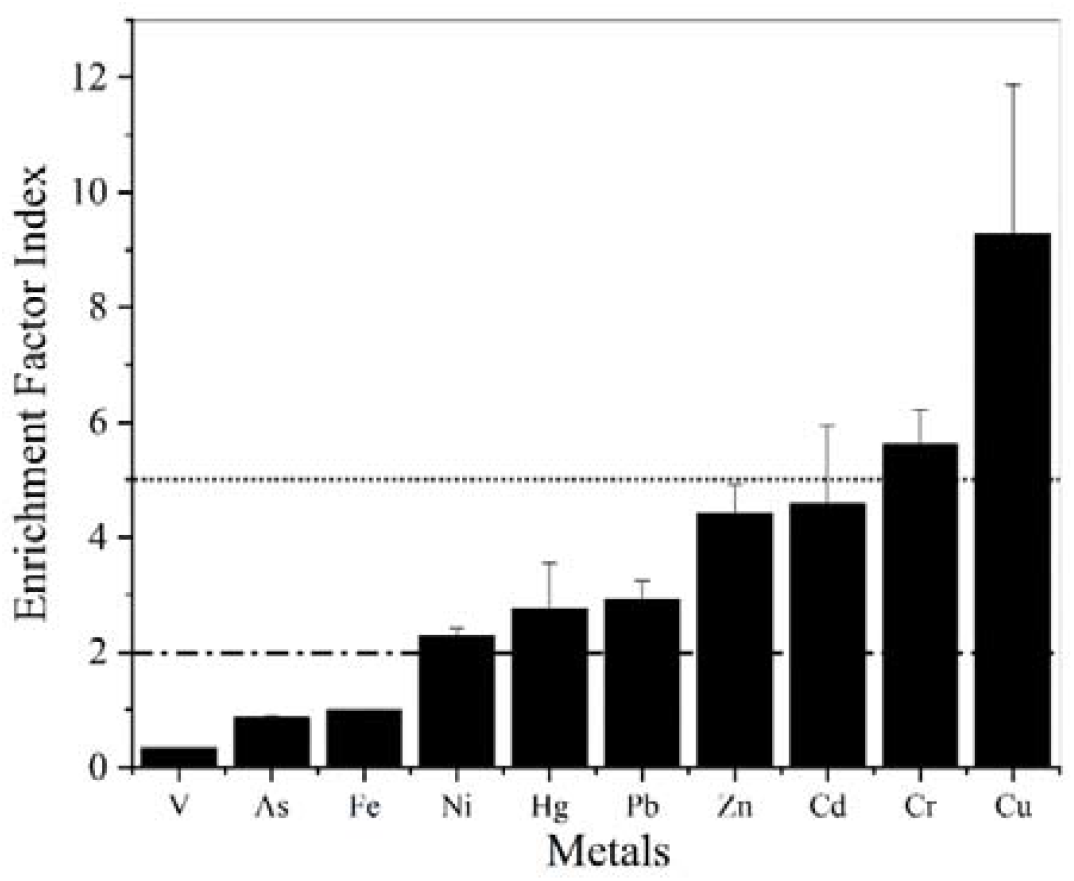
Enrichment factors (EFs) of heavy metals in Hong Kong’s marine sediments. Note: Values were normalized using Fe and local background levels in Hong Kong (Lau et al., 1993) and the earth crust average as references (Wedepohl, 1995). Error bar represents standard deviation.

### 3.3 Potential contamination index

The potential ecological risk indices 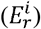 and *RI* for each site were obtained and listed in Table 3. According to these data, As, Cr, Pb, Ni, Zn, Fe and V posed a low ecological risk in all areas. However, the ecological risk for Cd in all regions showed moderate levels to severe risk levels (except the PS showed la ow level). Additionally, Cu posed a medium ecological risk in JS and ES, and a considerable ecological risk in VS. Other areas exhibited a low ecological risk. In comparing, Hg posed a low ecological risk in MS and TS, moderate ecological risk in ES, a and a high ecological risk in VS and JS. The rest of the areas displayed low ecological risk. In summary, the potential ecological risk indices for single metals 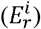 indicate that the severity of pollution of the ten heavy metals decreased in the following sequence: Cd > Hg > Cu > Pb > Ni > Cr > As > Zn > Fe > V.

**Table3.** Heavy metal potential ecological risk indexes of the Hong Kong coastal areas.

| Site | $E_r^i$ | | | | | | | | | | RI |
| --- | --- | --- | --- | --- | --- | --- | --- | --- | --- | --- | --- |
|  | As | Cd | Cr | Cu | Pb | Hg | Ni | Zn | Fe | V |  |
| DS | 13.17 | **168.75** | 12.04 | 37.68 | 14.64 | **70.36** | 15.68 | 4.81 | 1.17 | 0.78 | 339.07 |
| NS | 10.31 | **81.00** | 8.94 | 23.07 | 11.74 | **54.86** | 12.18 | 3.33 | 1.05 | 0.69 | 207.17 |
| SS | 7.39 | **43.13** | 9.39 | 24.91 | 10.33 | **63.39** | 12.49 | 2.99 | 0.95 | 0.64 | 175.61 |
| WS | 7.90 | **45.00** | 9.07 | 23.57 | 10.00 | **70.00** | 11.55 | 3.65 | 0.96 | 0.59 | 182.29 |
| VS | 8.28 | **387.50** | 19.56 | **141.01** | 19.25 | **272.62** | 18.53 | 7.13 | 0.89 | 0.61 | 875.38 |
| JS | 5.88 | **45.00** | 10.00 | **43.93** | 10.39 | **211.43** | 11.40 | 3.04 | 0.78 | 0.52 | 342.37 |
| ES | 5.78 | **117.00** | 10.91 | **61.86** | 13.18 | **118.29** | 10.86 | 4.31 | 0.82 | 0.50 | 343.51 |
| MS | 6.83 | **58.75** | 9.08 | 18.54 | 10.61 | 24.52 | 14.08 | 3.15 | 1.02 | 0.69 | 147.28 |
| TS | 8.61 | **276.00** | 8.17 | 35.57 | 23.26 | 30.57 | 11.82 | 5.78 | 0.94 | 0.71 | 401.43 |
| PS | 5.82 | 39.00 | 6.49 | 16.57 | 9.24 | **44.86** | 8.22 | 2.49 | 0.71 | 0.44 | 133.83 |
| AVE | 8.00 | **126.11** | 10.37 | **42.67** | 13.27 | **96.09** | 12.68 | 4.07 | 0.93 | 0.62 | 314.79 |

Notes: $E_r^i$ is the individual heavy metal potential ecological risk index, $RI$ is the total heavy metal potential ecological risk index; bold types indicate the heavy metals in sample sites with having ecological risk.

Furthermore, the spatial distribution of the potential ecological risk indices for all factors (*RI*) is shown in Fig. 4. The areas such as SS, WS, MS and PS showed moderate potential ecological risk, which indicates that industrial or anthropogenic activities have caused heavy metal pollutions and ecological risks in these areas. By contrast, considerable potential ecological risks were found in DS, NS, JS and ES, which implies potentially higher ecological risks and more severe environmental pollution than that in above areas. It is worth noting the severe potential ecological risks were characterized in VS and TS, which suggests that heavy metals have caused significant contamination in the sediments of the coastal region.

**Fig. 4.**
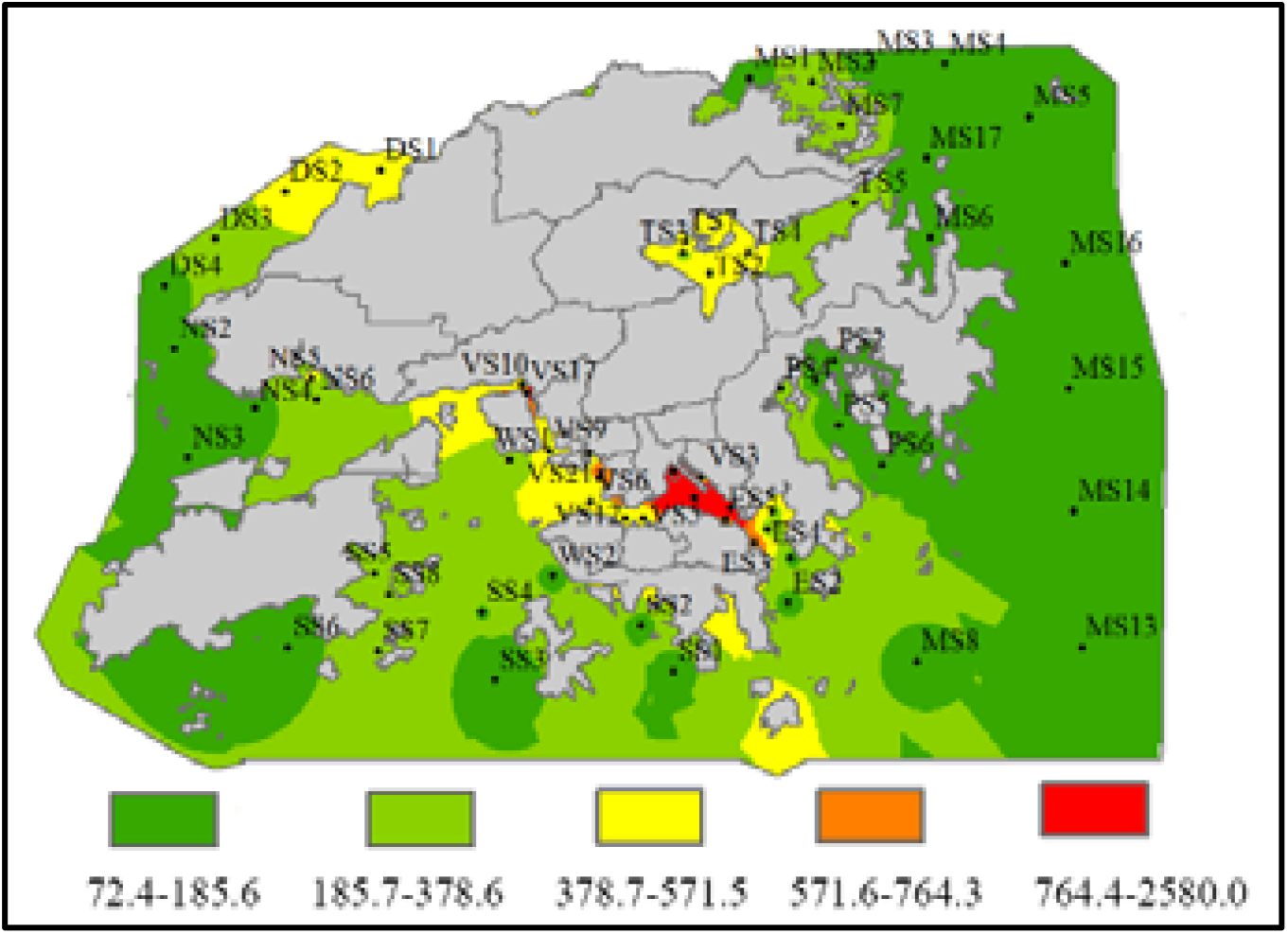
Spatial distribution of potential ecological risk indices (RI).

**Fig.5.**
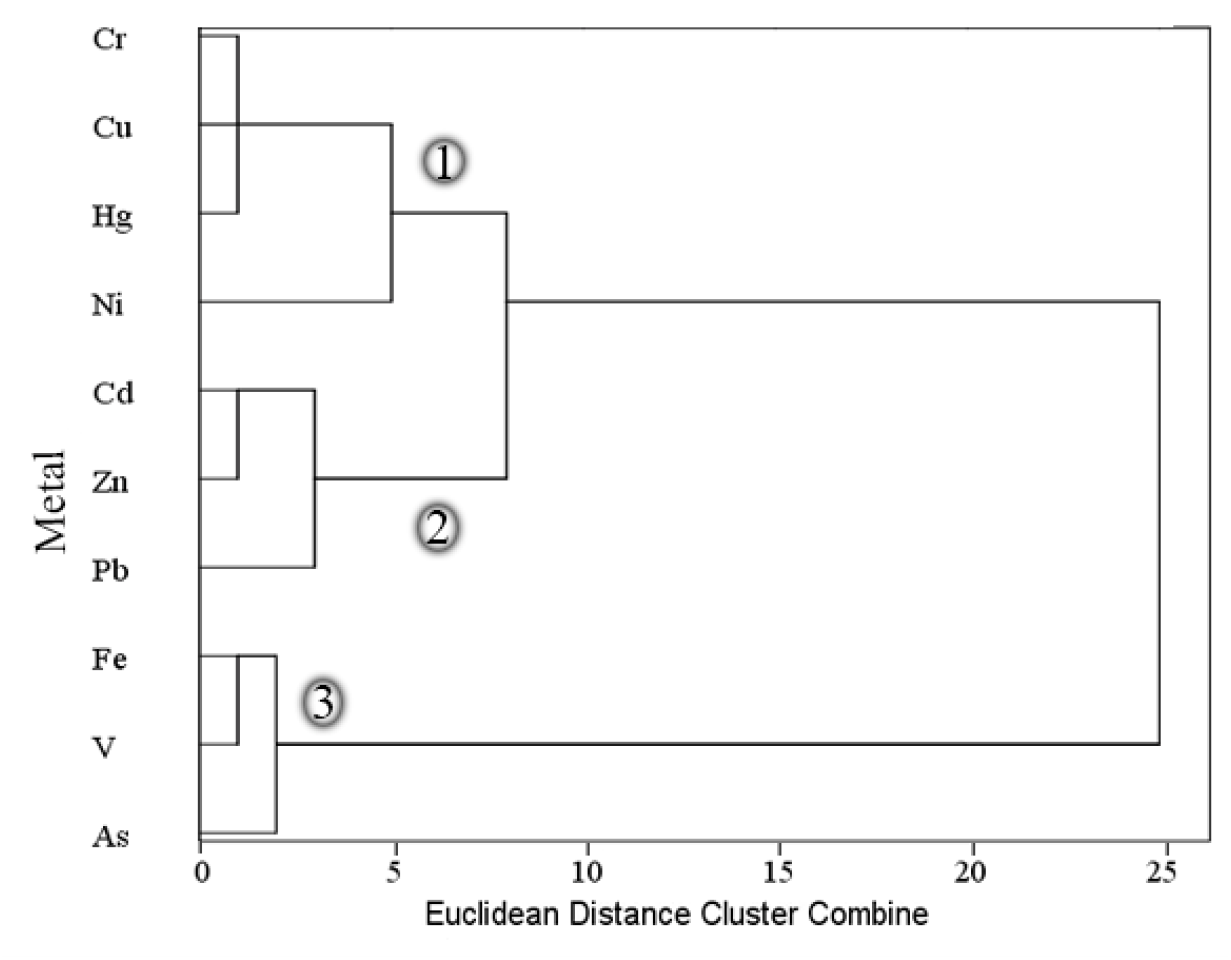
Dendrogram of cluster analysis for variables in the sediments of the study area.

**Fig.6.**
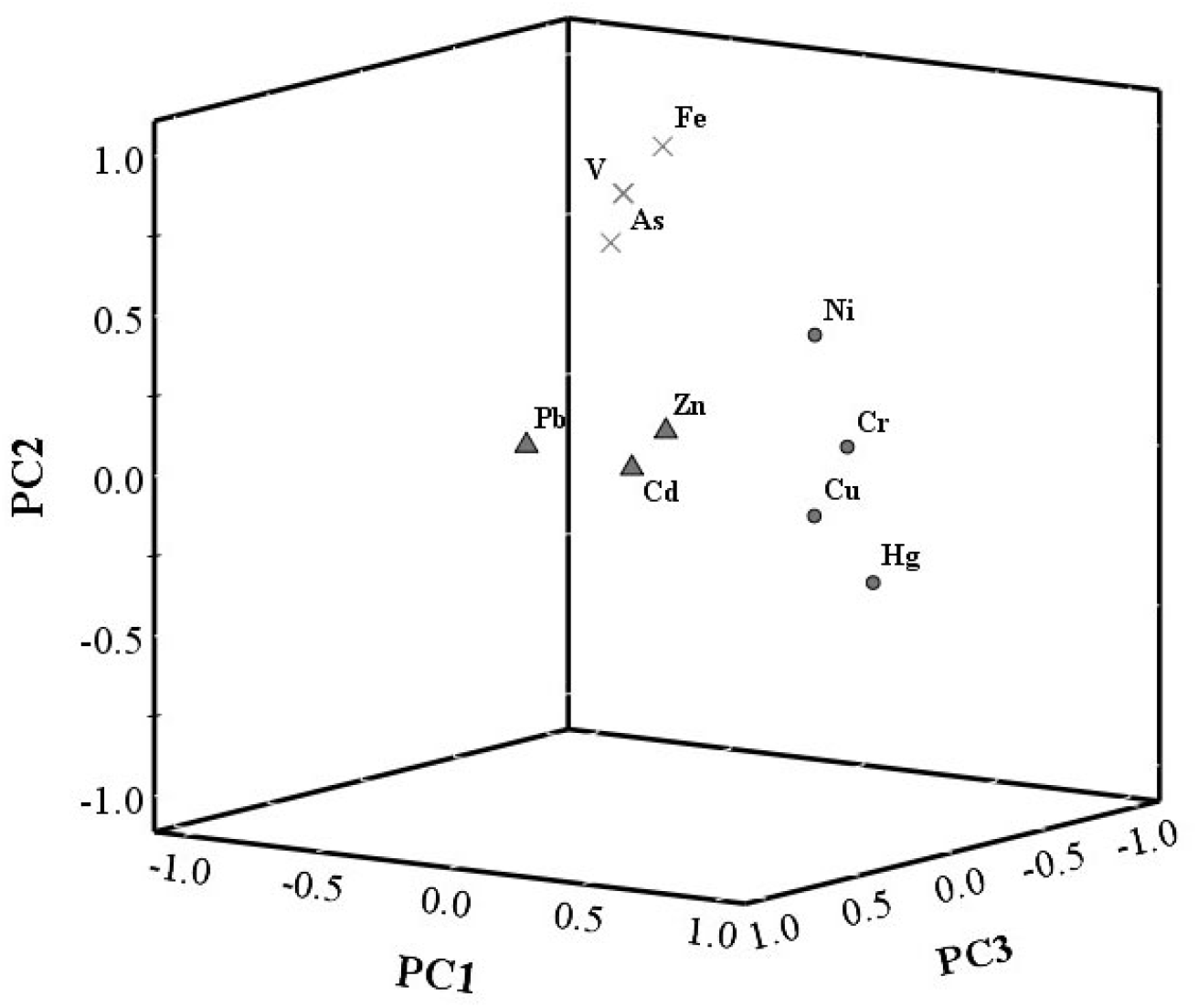
A 3D plot of scores obtained from PCA results for variables in the sediments of the study area.

### 3.4 Multivariate statistical analyses

Different methods commonly used for distinguishing between geogenic and anthropogenic sources of the potentially toxic elements include element speciation, profile distribution and spatial distribution. These methods are not accurate enough to distinguish between sources of element concentration on their own and should be combined with additional parameters such as a parent rock composition or known anthropogenic loads (Chandrasekaran et al., 2015). Multivariable statistical analysis provides an alternative method to recognize pollution sources and apportion geogenic versus anthropogenic contributions (Zhang et al., 2020). Principal component analysis (PCA) and its derivative methods have been widely used to achieve the objective. Cluster analysis (CA) is often coupled to PCA to check results and provide grouping of individual parameters and variables. PCA in conjunction with CA provides a means for ensuring proper source identification for a given metal distribution pattern in sediment (XXX).

#### Cluster analysis (CA)

The elements were standardized employing z scores prior to CA, and Euclidean distances for similarities in the elements were calculated. Finally, the hierarchical clustering by applying Ward’s method was performed on the standardized data set (XXX). All ten elements were grouped into two statistically meaningful clusters based on Euclidean distances. The constructed dendrogram showed two major distinct clusters with three groups formed. Cr, Cu, Hg and Ni have good similarities and were clustered in one group, whereas Cd, Zn and Pb were clustered in another group. These two groups showed a close relationship in one of the major clusters, indicating strong anthropogenic sources. Fe, V and As were clustered into the third group, while no close similarity is shown with the rest of the group. The third group formed the second cluster, which can mainly be originated from natural sources, because Fe is mainly derived from lithogenic sources.

#### Principal component analysis PCA

PCA was performed to identify the different sources of heavy metals in the study area. Three significant principal components and their percentage of the total variance were determined and shown in Table 4. At first, to examine the suitability of data for PCA, KMO and Bartlett’s Sphericity tests were accomplished. The calculated value of KMO is 0.856 and the significance level of Bartlett’s Sphericity is 0 (less than 0.05), which generally indicates the compatibility of data for PCA/FA. Therefore, this statistical analysis can be useful.

**Table 4.** The result of PCA for the entire data set.

| Element | PC1 | PC2 | PC3 |
| --- | --- | --- | --- |
| Cd | 0.49 | 0.15 | **0.83** |
| Cr | **0.93** | 0.18 | 0.31 |
| Cu | **0.89** | -0.02 | 0.43 |
| Pb | 0.48 | 0.20 | **0.96** |
| Hg | **0.90** | -0.27 | 0.13 |
| Ni | **0.78** | 0.51 | 0.27 |
| Zn | 0.56 | 0.26 | **0.75** |
| V | 0.08 | **0.89** | 0.30 |
| As | -0.02 | **0.86** | 0.15 |
| Fe | 0.04 | **0.99** | 0.02 |
| Eigenvalue | 3.35 | 3.29 | 2.93 |
| Total variance % | 36.84 | 30.05 | 26.55 |
| Cumulative % | 36.84 | 66.89 | 93.44 |

Note: Bold values indicate strong loadings.

PCA was applied to the normalized data set (10 variables) to identify the factors affecting each variable and also to develop the new pollution index in this study. The result of PCA demonstrated that three PCs with eigenvalues >1 were extracted which explain 93.44 % of total variance in respective data set and are presented in a three-dimensional space. The values of eigenvalue specify the significance of the PCs. Eigenvalues bigger than 1 are considered significant. The obtained values of factor analysis including loadings, eigenvalues and %variance are presented in Table 4. The coefficients greater than 0.7 were considered as significant or strong loading values (Pejman et al., 2015).

PC1 accounts for 36.84 % of the total variance and has strong positive loadings on Cr, Cu, Hg and Ni which can be interpreted as anthropogenic pollution sources. PC2 explains 30.05 % of the total variance and has strong positive loadings on V, As and Fe, representing common natural sources. PC3 explains 26.55 % of the total variance and has strong positive loadings on Cd, Pb and Zn. These elements also showed a strong positive correlation with PC1, indicating mainly affected by anthropogenic pollution sources.

#### Source identification of heavy metals

Hong Kong is a highly urbanized coastal city receiving substantial metal loads from industrial and municipal wastewaters (Xu et al., 2011). Metal contamination of marine sediment in Hong Kong through many pathways, including disposal of liquid effluents, terrestrial runoff and leachates carrying chemicals that originated from numerous urban, industrial wastes and shipping activities, also atmospheric deposition (Holmes, 1996; Ruan et al., 2019; Jing et al., 2020). It is worth noticing that the rapid industrialization and urbanization in the Pearl River Delta area of south China during the last few decades have led to widespread contamination of heavy metals in the coastal and estuarine sediments of Hong Kong, especially in the Deep Bay and North Southern Bay, a semi-enclosed bay that close to Shenzhen Bay (Tian et al., 2014; Duan et al., 2021). It receives discharges from the Shenzhen river all year round. Moreover, the water quality in two areas has also been influenced by the Pearl River during the wet season (Hong et al., 2010). Therefore, the sediment quality in these areas might be significantly affected by the discharge from Shenzhen River and Pearl River, especially for some heavy metals and organic pollutants. Victoria Harbour and adjacent areas (Western Buffer, Junk Bay and Eastern Buff) are major ports of Hong Kong city and locate between the most urbanized areas of the Kowloon Peninsula and the northern shore of Hong Kong Island. These regions have been under tremendous pressure of environmental pollution, due to the increasing discharges of industrial effluents, domestic sewage via runoffs along the bank (Nicholson et al., 2011; Archana et al., 2016). The Southern Bay and Mirs Bay, which lie at the southern and southeast corner of Hong Kong in the open sea, are relatively far from local islands, thus less influenced by urbanization and industrialization. The eastern offshore areas (Tolo Harbour and Port Shelter) experienced serious pollutions due to excessive sewage loadings, including the industrial effluent discharges and wastewaters following the development of new towns around its shores (Lai et al., 2016).

From the statistical analyses, three main sources of heavy metals in sediments from the coast of Hong Kong have been identified. The first principal component (PC1) consists of four metals: Cr, Cu, Hg and Ni. Cr occurs naturally in the earth’s crust, but its extensive use in various industrial processes and products has led to widespread chromium contamination in the environment (Guan et al., 2014). High concentrations of Cr due to the disposal of wastewater from industries such as metal fabrication and chromium plating (Coetzee et al., 2020). Additional areas of application of chromium include wood preservatives, chrome pigments used in paints, printing inks and anti-corrosive materials for ship constructions (Balgude et al., 2012). All these metal industries may contribute Cr to the sediment pollution in the Hong Kong coastal area, especially during the past decades with the rapid tourism and trade-transportation development in Hongkong. Cu is a major marine toxicant of concern owing to its widespread use in the shipping industries. The use of copper as the primary active ingredient in antifouling paints contributes a significant portion of copper influx into the marine environment (Cerchier et al., 2020). Both are common industries in Hong Kong city. Industrial and urban centers along coastal areas are major sources of Hg pollution in oceans (Liu et al., 2020). Another major source of atmospheric Hg emission is the combustion of coal in coal-fired power plants. Hong Kong contains eight coal-fired stations operated by the Hong Kong Electric Company (HEC) and several large industrial estates which are typically situated near important water systems (Moss et al., 2016). Ni has been major metals pollutions in Hong Kong area due to the various pollution inputs from many industrial processing factories, combustion of marine fuel in ships and sewage sludge from sewage treatment plants according to the literature (Li et al., 2017). Taken together, the high concentrations of Cr, Cu, Hg and Ni in PC 1 are closely related to industrial inputs and shipping activities around Hong Kong city.

The second principal component (PC2) contains three metals: V, As and Fe. V is widely distributed in igneous and sedimentary rocks and minerals as a mildly incompatible, refractory, lithophilic element (XXX). Similarly, As is primarily associated with igneous and sedimentary rocks, which are mobilized naturally through volcanic, geothermal and microbiological processes and by weathering of crustal rocks (Peshut et al. 2008). Besides, Fe in the sediment originates largely from the earth’s crust or natural rock via weathering (XXX). In comparison to PC 1 and PC 3, the mean concentration of these elements in PC2 approximates or not exceeds its background value, indicating they are mainly derived from natural origin rather than anthropogenic contamination.

The third principal component (PC3) includes three metals: Cd, Pb and Zn. Cd is one of the most important toxic metals in industrial discharges. A major source of Cd pollution in the environment comes from industrial wastewater steam, such as effluents from electroplating, battery, metal finishing and printed circuit board industries (Krishnan et al., 2021). Similarly, the important sources of Pb include industrial discharges and the printed circuit industry (Wang et al., 2015). In Hong Kong, a gradual reduction in the amount of Pb permitted in leaded petrol began in the early 1980s, and leaded petrol was totally banned in 1999. including the leaded gasoline used in the past (Wang et al., 2011). Consequently, the increased enrichment of Pb most likely originated from the discharges of industrial wastewater. Zn concentration is higher than its background values. It was apparently attributed to nearby municipal or industrial activities. In recent years, as water-transportation activities and related tourism were further intensified in the Hong Kong, the heavy metals from municipal waste increased significantly. Therefore, sources for this component (PC2) might be identified from industrial wastewater and municipal discharges.

## Conclusion

In this work, ten elements in sediments from the coastal area of Hong Kong were investigated. The spatial distributions of the ten elements have been analyzed, and their pollution degrees and corresponding potential ecological risks and the source identification have been studied using the enrichment factor analysis, potential ecological risk index and integrated multivariate statistical methods, respectively. Except V, Fe and As, the mean concentrations of all the other heavy metals in sediments exceed their geochemical background values. According to the results obtained from the enrichment factor analysis, Cu and Cr are identified as the primary metal pollutants. Based on potential ecological risk indices, severe potential ecological risks were characterized in VS and TS, which suggests that heavy metals have caused significant contamination in the sediments of the coastal region. Multivariate statistical analyses show that Cr, Cu, Hg and Ni may originate from industrial inputs and shipping activities, while V, As and Fe are identified from natural sources, and Cd, Pb and Zn are derived from industrial wastewater and municipal discharges. This work should be useful to the establishment of strategies for contamination control and management and the optimization of industrial structures in Hong Kong.

## Acknowledgements

This research received no specific grant from any funding agency in the public, commercial, or not-for-profit sectors.

## Author contributions

Conceived and designed the study: C Chen, FW Huang; analyzed the data: C Chen, FW Huang; drafted the manuscript: C Chen, FW Huang.

